# Engineering CRBN for rapid identification of next generation binders

**DOI:** 10.1101/2024.01.18.576231

**Authors:** Henry J. Bailey, Jonathan Eisert, Joshua Vollrath, Eva-Maria Leibrock, Ivan Kondratov, Tetiana Matviyuk, Nataliya Tolmachova, Julian D. Langer, Ansgar A. Wegener, Fiona J. Sorrell, Ivan Dikic

## Abstract

The majority of clinical degrader candidates utilize an immunomodulatory imide drug (IMiD)-based derivative, that directs their target substrate to the E3 ligase receptor Cereblon (CRBN), however, concomitant neo-substrate degradation by IMiDs often results in severe off-target effects. Further biophysical screening is needed to discover CRBN binders that might overcome these safety concerns, but the previously reported CRBN protein constructs suffer significant limitations that reduce their applicability as tools for biophysical assays making large-scale screening efforts a costly endeavor. This is due to the requirement of CRBN co-expression with DDB1 in insect cells to produce soluble protein that contains all the reported structural features necessary for proper compound interaction. Here, a near full-length human CRBN construct was designed that retains these required features, but for the first time allows the generation of highly homogenous and cost-efficient expression in *E.coli*, eliminating the need for DDB1 co-expression. We have extensively profiled the new construct against the existing CRBN constructs in binary and ternary degrader complexes, showing it to be the “best of both worlds” in terms of binding characteristics and ease of protein production. We then designed the Enamine CRBN focused IMiD library of 4480-compounds and demonstrated applicability of the new construct to high throughput screening by identifying novel CRBN binders with high potency and ligand efficiency. The new methods described here should serve as valuable tools for the development of the next generation IMiD-based degraders.

## Introduction

Targeted protein degradation (TPD) has emerged as an important therapeutic modality that utilizes bifunctional degrader molecules, such as proteolysis targeting chimeras (PROTACs) or molecular glues (MGs), to recognize and recruit specific proteins to E3 ligase complexes for ubiquitination and subsequent degradation via the cellular ubiquitin-proteasome system^1–3^. The majority of PROTACs and MGs currently in clinical trials utilize an IMiD-based derivative, such as thalidomide, lenalidomide or pomalidomide, that directs their target substrate to the E3 ligase receptor Cereblon (CRBN), making CRBN the most successful effector protein within the TPD field^4–6^.

CRBN has a modular structure (Figure 1A) with an N-terminal Lon protease-like domain, an intermedial helical bundle domain (HBD) followed by a C-terminal thalidomide binding domain (TBD) that binds the glutarimide moiety of IMiDs within a tryptophan cage pocket ^7–10^. In addition to the ligand binding pocket, the TBD encodes a sensor loop that allows for conformational crosstalk between the TBD and Lon domain during compound binding^11^. Second generation high affinity elaborations of the core IMiD scaffold, such as mezigdomide and iberdomide, harness this crosstalk through direct interactions with the sensor loop, favoring transition to a closed CRBN conformation that allows the Lon N domain to provide stabilizing interactions to the TBD pocket^11^. In contrast, the helical bundle domain does not participate in ligand binding, instead it contributes 55 amino acids which are responsible for recruiting the CUL4 RING ubiquitin ligase machinery via direct interaction with the adapter protein DDB1(Figure 1A)^8,10^.

**Figure 1.**
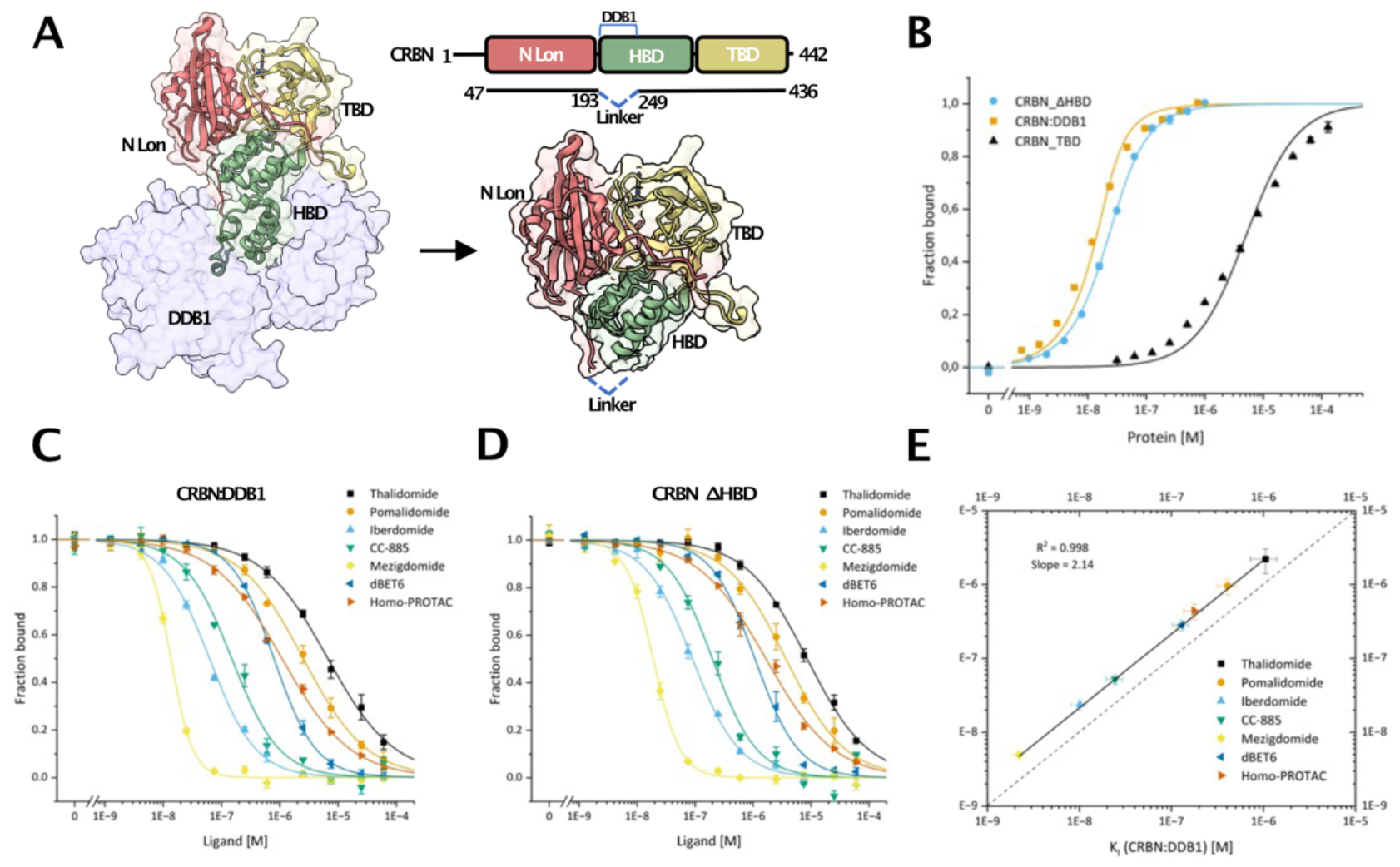
New CRBN construct retains binary tool compound affinity: **A)** Linker design for expression and purification of soluble near full-length CRBN (CRBN_ΔHBD) in *E.coli*. **B)** Fluorescence polarization assay comparing affinity for thalidomide-based Cy5 tracer to CRBN_ΔHBD versus the minimal thalidomide binding domain (TBD) and CRBN:DDB1 complex **C)** Competitive fluorescence polarization assay comparing K_i_ of published tool CRBN PROTACs and molecular glues against CRBN:DDB1 complex. **D)** Competitive fluorescence polarization assay comparing K_i_ of published tool CRBN PROTACs and molecular glues against CRBN_ΔHBD. **E)** K_i_ correlations of tool CRBN compounds between CRBN_ΔHBD and the CRBN:DDB1 complex.

Two recombinant human CRBN constructs have been previously described for characterization of CRBN-based degraders, however, significant hurdles exist that limit their applicability for biophysical screening methods. The minimal TBD (CRBN_TBD) construct is expressed in *E. coli* cells but lacks important contributions from the Lon N domain and sensor loop that limit its activity and affinity during IMiD binding^11,12^. A bacterial cereblon isoform from *Magnetospirillum gryphiswaldense* (MsCI4), designed to correspond to the human TBD region for high throughput ligand screening, is also shown to bind ligands with affinities several orders of magnitude lower than reported for human full-length cereblon^13^. A second near full-length human CRBN construct exists that overcomes these technical issues, however, to retain CRBN solubility it requires specialized and costly co-expression in insect cells as a heterodimer with the interaction partner DDB1^8^ and is heterogeneous due to dissociation of the complex, presenting challenges for structural and biophysical assays.

To tackle this we engineer an intermediate human CRBN construct (CRBN_ΔHBD), that contains all important regions for compound binding, and purify from a simple, convenient *E.coli* expression system. We develop several binary and ternary interaction assays and show highly desirable properties of CRBN_ΔHBD for the evaluation of PROTACs and molecular glues providing invaluable tools to delineate structure activity relationships of IMiD based degrader compounds. Finally, we optimize high-throughput ligand primary screening assays and couple these with a focused CRBN library supplied by Enamine to discover next generation CRBN binders with improved binary affinity and ligand efficiency, indicating this new construct will be valuable in the search for new CRBN binders and alternative scaffolds.

## Results

We hypothesized the limiting factor preventing soluble full-length human CRBN production is exposure of several hydrophobic patches in the HBD to solution in the absence of the binding partner DDB1. Inspection of the CRBN:DDB1 complex structure^8,10^ revealed the DDB1 interacting residues (aa 194-248) of the HBD could easily be deleted and replaced with a soluble GNGNSG linker to form an intermediate CRBN construct (CRBN_ΔHBD) (Figure 1A). AlphaFold models show CRBN_ΔHBD can adopt the functional closed conformation with an RMSD of 0.6 across 309 residues when overlayed with PDB:6BOY^14^ indicating deletion of the DDB1 interacting sequence does not perturb the structure of the TBD and Lon N domains (Supplementary figure 1A). MBP-His-CRBN_ΔHBD was expressed in *E. coli* and isolated in a three-step purification process using IMAC, tag cleavage, SEC and IEX to greater than 95 percent purity (Supplementary figure 1B). To show CRBN_ΔHBD is a suitable construct for studying binary ligand interactions, affinity differences between purified CRBN:DDB1, CRBN_TBD and CRBN_ΔHBD were measured against a thalidomide-derived Cy5 tracer molecule in fluorescence polarization (FP) assays^14^ (Figure 1B)(Supplementary Table 1). Similar to previous studies^12^, isolated CRBN_TBD has significantly lower affinity towards the tracer (5000 nM) when compared with the near full-length CRBN:DDB1 heterodimer (4.1 nM). However, the newly engineered construct, CRBN_ΔHBD, binds the tracer molecule with a K_D_ of 13 nM, showing highly similar affinity to CRBN:DDB1. To confirm this behavior across a range of first- and second-generation MGs and PROTACs (Supplementary Table 1)(^14–19^), competitive FP experiments were performed resulting in highly comparable K_i_ values (R^2^ values of 0.998) between CRBN:DDB1 and CRBN_ΔHBD constructs (Figure 1C,D,E). This indicates that CRBN_ΔHBD, like CRBN:DDB1, is able to adopt a fully functional conformation where engagement of the sensor loop and Lon N domains is required for interaction with high affinity second-generation MGs such as mezigdomide^11^. Stability, and activity of CRBN_ΔHBD was further validated with thermal shift measurements showing monophasic melting curves with mid-point shifts between 6 and 14 °C in the presence of the ligands (Supplementary figure 2A), correlating well with measured K_i_ (Supplementary figure 2B).

To investigate CRBN_ΔHBD activity in ternary complex formation assays the bromodomain-containing protein 4 (BRD4) PROTAC dBET6^14^ was selected (Supplementary table 1). Spectral shift assays were optimized to directly calculate ternary K_D_ and apparent cooperativity value (alpha) in the presence of the first bromodomain of purified BRD4 (BRD4-BD1, amino acids 42-168) (Figure 2A). In line with previously reported literature values dBET6 displays a negative apparent alpha cooperativity of 0.5^14^. For direct visualization of ternary complex formation, SEC-HPLC elution profiles were recorded for both CRBN_ΔHBD and BRD4-BD1 in presence of dBET6 and additionally CRBN_ΔHBD in the presence of the CRBN homo-PROTAC^19^ (Supplementary Figure 3). Both PROTACs yield almost 100 percent ternary complex formation seen through a reduction in retention volume of the individual monomer peaks to single ternary complex elution with increasing PROTAC concentration (Supplementary Figure 3). Ternary complex formation between BRD4-BD1 and CRBN_ΔHBD was further validated in the presence of both dBET6 and the covalent lysine crosslinker Disuccinimidyl suberate (DSS, Supplementary table 1) using denaturing SDS-PAGE (Figure 2B). Four inter-crosslinked peptides were identified by mass spectrometry (Figure 2B) and mapped onto the crystal structure of CRBN:DDB1-BRD4-dBET6 (PDB:6BOY) (Figure 2B)^14^. When compared against the structure, one of these crosslinks formed with an optimal distance of 30 Å between lysine Cα atoms, indicating the ternary complex with CRBN_ΔHBD is able to adopt a similar conformation to the single state resolved by X-ray crystallography. Three further inter-crosslinks formed with distance restraints greater than the optimal 30 Å, indicating the BRD4-CRBN_ΔHBD ternary complex may sample multiple conformations in solution. Rotation of BRD4-BD1 along a pivot point within the flexible dBET6 linker allowed for conformations with appropriate distance restraints for crosslinking (Figure 2B). The dynamic nature of the BRD4:CRBN ternary complexes has been previously reported using HDX mass spectrometry^20^, thus providing further evidence here that single snapshots of ternary complexes via X-ray crystallography are not representative of the ensemble of solution states.

**Figure 2.**
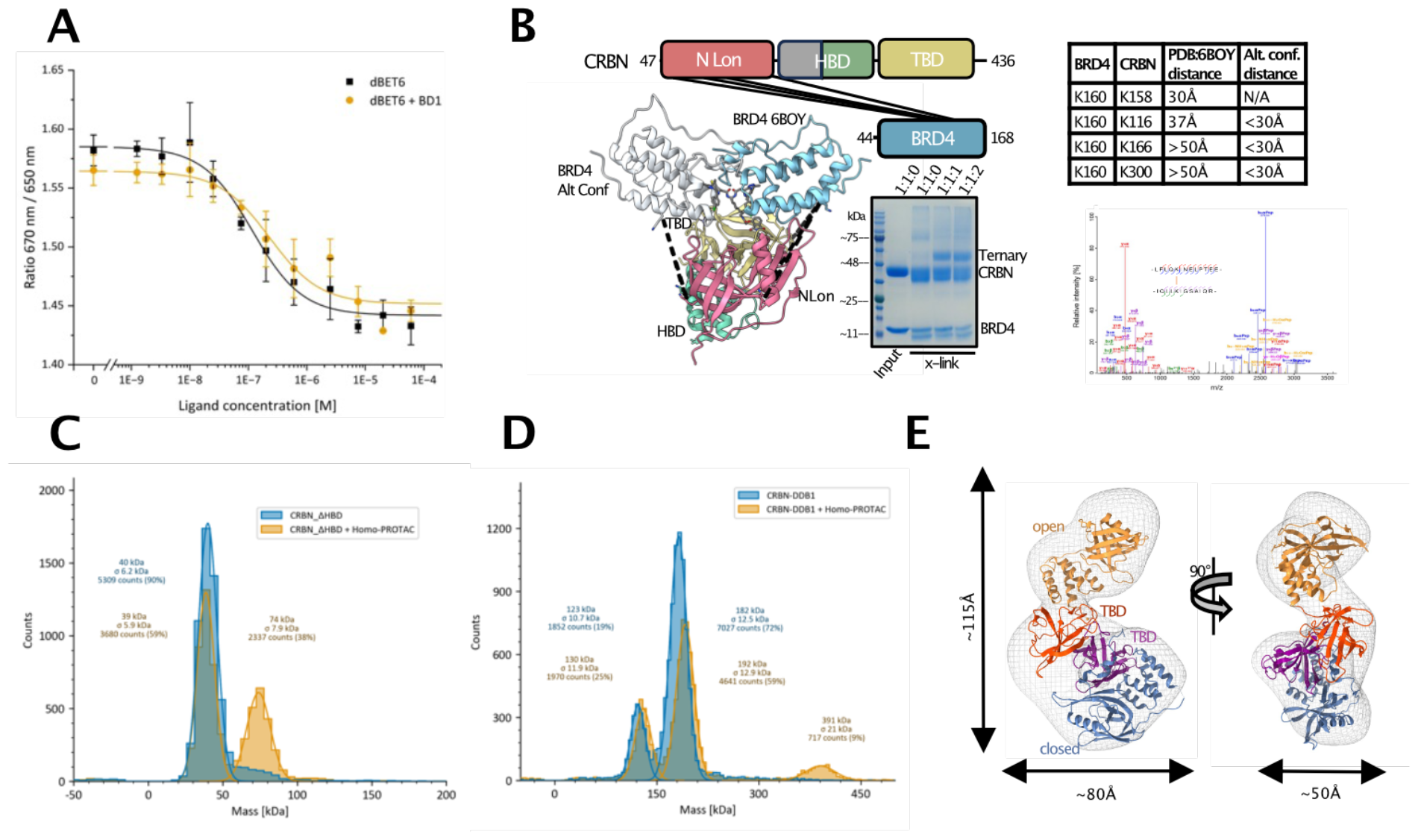
Visualization of complex formation with published tool PROTACs: **A)** Spectral shift affinity and cooperativity measurements of binary and ternary complex formation with CRBN_ΔHBD and BRD4-BD1 **B)** SDS-PAGE gel showing covalent capture of a BRD4-CRBN_ΔHBD ternary complex in the presence of lysine crosslinker and dBET6. Lane 1 contains protein MW marker, Lane two contains the input BRD4-BD1 and CRBN_ΔHBD at 1:1 molar ratio without DSS crosslinker, lane 3-5 contains 0.2 mM DSS crosslinker with a molar ratio BRD4-BD1:CRBN_ΔHBD:dBET6 of 1:1:0 (lane 3) 1:1:1 (lane 4) and 1:1:2 (lane 5). Crosslinked lysines within the BRD4-CRBN_ΔHBD dBET6 induced ternary complex identified via LC-MS/MS mapped onto the polypeptide scheme and crystal structure of the BRD4-CRBN-dBET6 complex PDB 6BOY. BRD4 is colored blue in the crystal structure and grey in the alternative modelled state. The table shows the distances between C-α atoms of crosslinked lysines in both conformational states. Representative fragmentation spectrum of accepted crosslinks (bottom right). **C)** Refeyn mass photometry measurements of CRBN_ΔHBD showing direct visualization of ternary complex formation after incubation with homo-PROTAC **D)** Refeyn mass photometry measurements of CRBN:DDB1 complex showing direct visualization of ternary complex formation after incubation with homo-PROTAC. **E)** 3D envelopes calculated from negative stain electron microscopy of CRBN_ΔHBD ternary complex after incubation with homo-PROTAC. CRBN in closed conformation (PDB:6BOY) and open conformation (PDB:6BNB) were rigid-body fit into density.

Oligomeric homogeneity of CRBN_ΔHBD was confirmed by mass photometry (Refeyn) (Figure 2C), where a single homogenous monomer peak was seen at the expected molecular weight of 40 kDa. Following addition of a CRBN homo-PROTAC, successful ternary complex formation was apparent by appearance of a second peak corresponding to the expected mass of the ternary complex (Figure 2C). Direct comparison of CRBN_ΔHBD and CRBN:DDB1 complex using mass photometry highlights the less favorable, heterogenous nature of the CRBN:DDB1 complex (Figure 2D). Two peaks were evident in the *apo* sample, one corresponding to the expected mass of DDB1 alone at 120 kDa, and one corresponding to the CRBN:DDB1 hetero-complex. Addition of homo-PROTAC to the CRBN:DDB1 complex did not result in any change in the peak at 120 kDa, further indicating this peak corresponds to DDB1 alone (Figure 2D). The significant portion of un-complexed DDB1 is likely caused by the instability and dissociation of the CRBN:DDB1 heterodimer.

CRBN_ΔHBD ternary complex formation in the presence of homo-PROTAC was further interrogated by negative stain electron microscopy. The micrographs revealed highly homogenous monodisperse particles and clear 2D classes could be resolved (Supplementary Fig 4). Calculation of an electron density map at >15 Å and rigid body fit of both open and closed states of CRBN structures showed that one CRBN molecule is able to adopt an open state conformation with the opposing CRBN molecule adopting the closed state (Figure 2E).

Due to its small size and oral availability, thalidomide-based derivatives are highly favorable E3 handles for PROTACs. However, significant issues with off-target toxicity and stability of this chemical scaffold are well reported^21,22^. Having established CRBN_ΔHBD as a relevant construct to study binary and ternary complex formation using tool compounds, we next optimized high-throughput screening assays to search for the next generation of scaffolds that may overcome the intrinsic issues with thalidomide. FP assays were miniaturized to 384-well plates and CRBN_ΔHBD was screened against a CRBN-focused library of 4480 compounds (Enamine). The library was specially designed for the CRBN-screening campaigns and investigation of new CRBN-based MGs. Thus, 3650 compounds contain the thalidomide core with various substitutions in the phthaloyl ring as well as the glutarimide ring (Supplementary Figure 5A) and an additional 830 compounds contained scaffold variation with close similarity to typical IMiD chemotypes (Supplementary Figure 5B). Synthesis of most of the compounds was performed at Enamine starting from available glutarimide containing Building Blocks by coupling with diverse counterparts using typical parallel chemistry procedures (Supplementary Figure 5C). The primary FP screen was performed at a 500 nM compound concentration in duplicates using lenalidomide and iberdomide controls to allow differentiation of FP response between low and high affinity binders (Figure 3A). In total 83% of compounds have detectable binding response against CRBN with a calculated assay Z-prime score of 0.85. 3% of compounds have a binding response equal to iberdomide or greater, 21% of compounds showed a binding response between lenalidomide and iberdoimide including the previously well-characterized MG pomalidomide, 23% of compounds had a binding response equal to lenalidomide and 36% compounds display a weaker binding response, including the well-characterized MG thalidomide (Figure 3A). 175 compounds were selected for dose-response measurements and revealed an excellent correlation between calculated EC_50_ and initial response in the primary screen, indicating single shot values are highly applicable for interpretation of structure activity relationships (Supplementary Figure 5).

**Figure 3.**
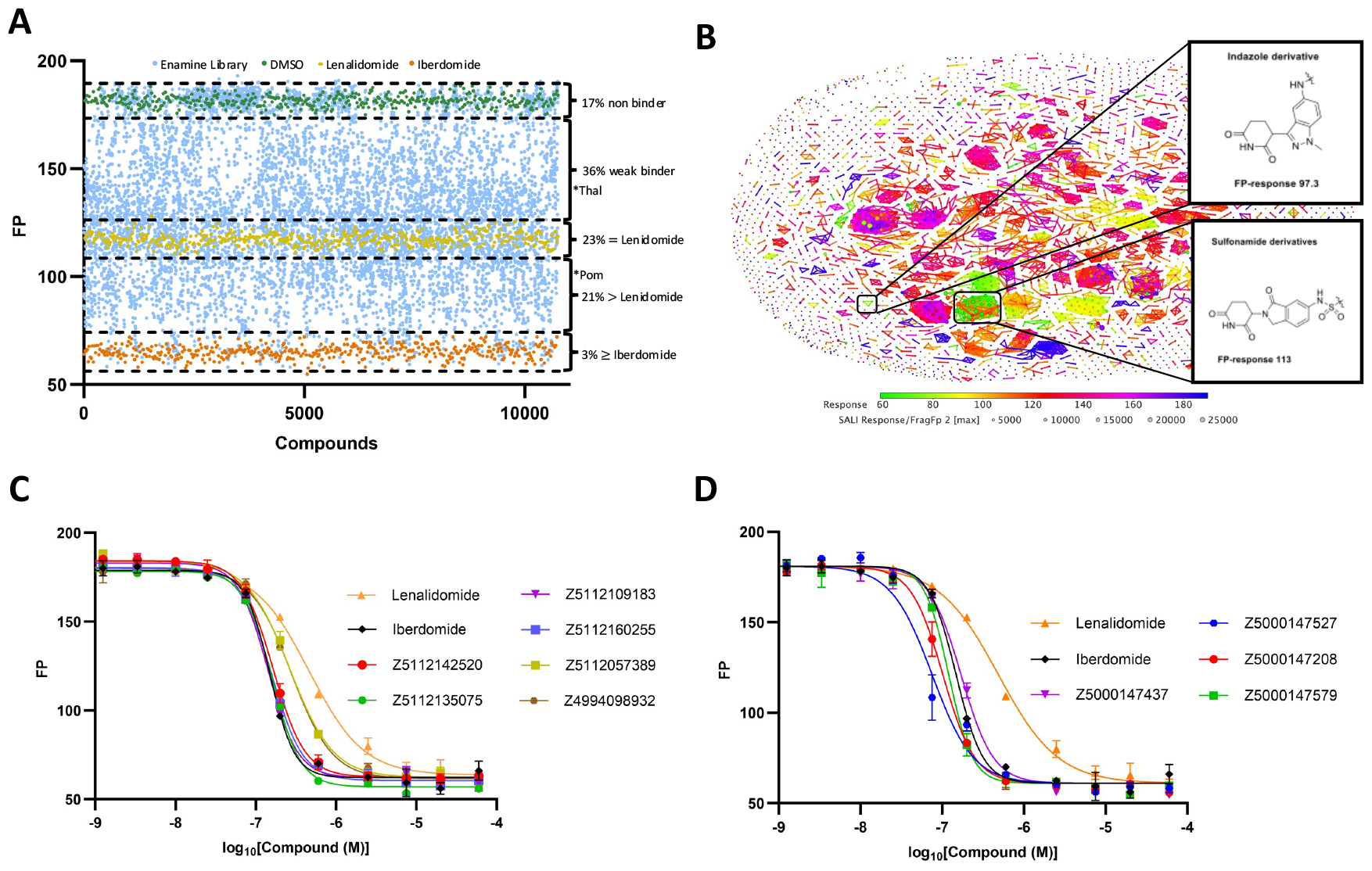
High-throughput screening for discovery of next generation CRBN binders: **A)** Single concentration fluorescence polarization competition screen of the focused CRBN library containing 4480 compounds measured in duplicate. Dashed lines represent 3 times standard deviation of control samples. Asterix shows response of tool compounds present in the Enamine library. **B)** Structure activity landscape plot of of FP response vs FragFP **C)** Dose response fluorescence polarization curves showing a subset of indazole based derivatives. **D)** Dose response fluorescence polarization curves showing a subset of sulfonamide-based IMiD derivatives.

Analysis of the structure activity landscape revealed that the high affinity compounds were predominantly derived from the sub-library of phthaloyl ring derivatives (Figure 3B). However, one indazole scaffold (Z4994098932), from the sub-library of IMiD scaffold variations, recently published to have affinity toward CRBN:DDB1 comparable to pomalidomide^23^, was also identified in this screen with very similar binding activity. (Figure 3B) (Table 1). Our data for this compound (Figure 3B,C) confirms the previously-reported findings, further validating this compound as a novel CRBN binder. Furthermore, 7 derivatives of this indazole scaffold present in the primary screen resulted in increased binding potency with EC_50_s in the range of iberdomide, with compound Z5112109183 showing most potent binding (Figure 3C) (Table 1). The indazole fragment has highly favorable trade-off between affinity, molecular weight and lipophilicity (clogP) values (Table 1) indicating its potential use in PROTACs which characteristically suffer from MW (>500 da) and lipophilicity values (clogP >5) beyond the classic rule-of-5^24^. All active compounds are derivatives of the core Building Block Z4994098932 with various substitutions in the amino group which is suitable as an exit vector for further derivatives (e.g. linker connection for PROTAC design). In addition to the active binders containing the indazole fragment, several weaker binders with new IMiD cores were identified (Supplementary Figure 7). Their further optimization can be used for development of potent CRBN-binders with new chemoptypes..

**Table 1:**
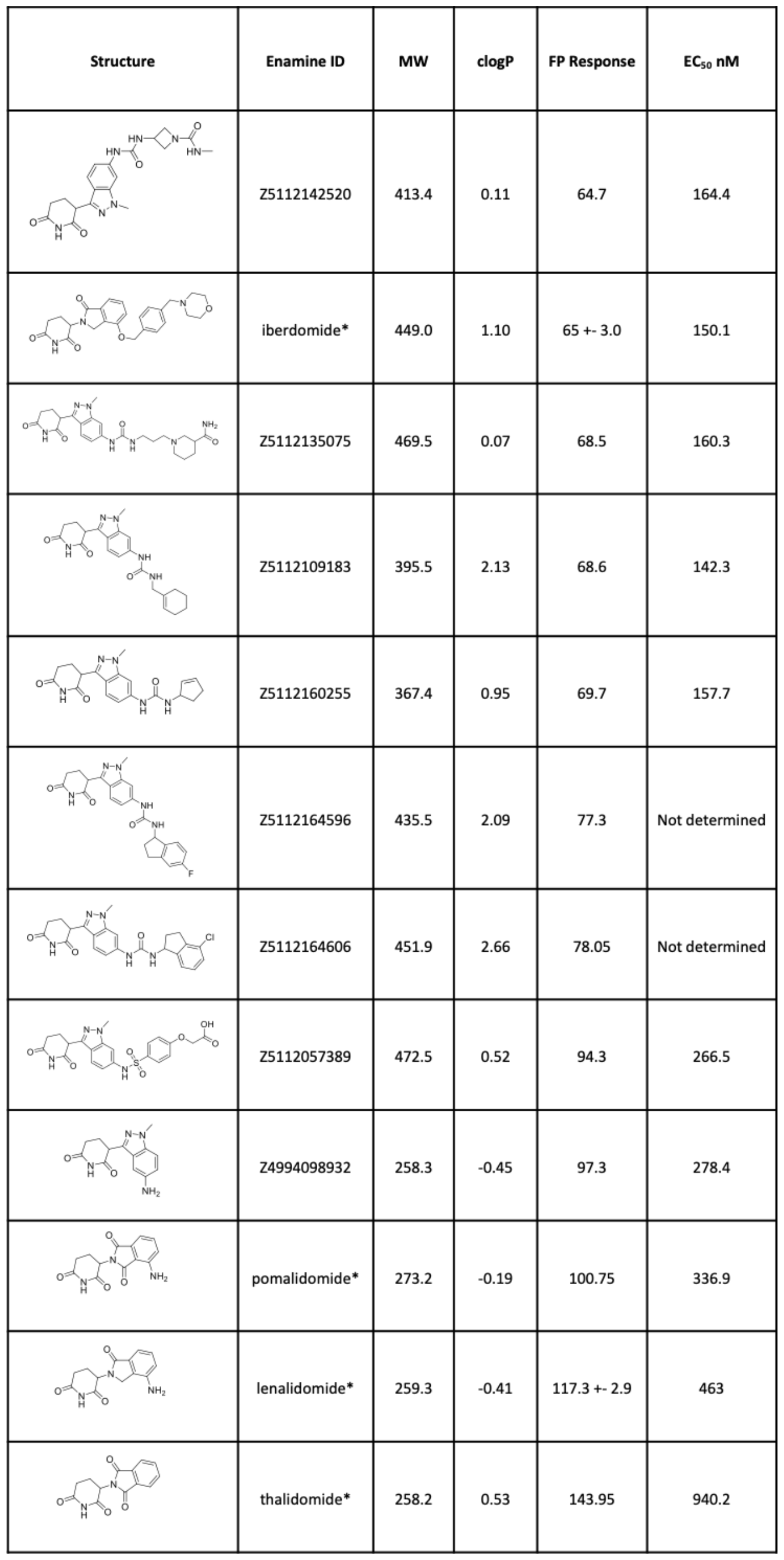
Indazole based CRBN binders. *=control compounds

| Structure | Enamine ID | MW | clogP | FP Response | EC<sub>50</sub> nM |
| --- | --- | --- | --- | --- | --- |
|  | Z5112142520 | 413.4 | 0.11 | 64.7 | 164.4 |
|  | iberdomide* | 449.0 | 1.10 | 65 +- 3.0 | 150.1 |
|  | Z5112135075 | 469.5 | 0.07 | 68.5 | 160.3 |
|  | Z5112109183 | 395.5 | 2.13 | 68.6 | 142.3 |
|  | Z5112160255 | 367.4 | 0.95 | 69.7 | 157.7 |
|  | Z5112164596 | 435.5 | 2.09 | 77.3 | Not determined |
|  | Z5112164606 | 451.9 | 2.66 | 78.05 | Not determined |
|  | Z5112057389 | 472.5 | 0.52 | 94.3 | 266.5 |
|  | Z4994098932 | 258.3 | -0.45 | 97.3 | 278.4 |
|  | pomalidomide* | 273.2 | -0.19 | 100.75 | 336.9 |
|  | lenalidomide* | 259.3 | -0.41 | 117.3 +- 2.9 | 463 |
|  | thalidomide* | 258.2 | 0.53 | 143.95 | 940.2 |

We next filtered the screening results based on FP response vs. molecular weight and FP response vs. clogP (Figure 4A,B) values and identified a further 7 potent scaffolds with EC_50_ values greater than pomalidomide (Figure 4C and Table 3) whilst retaining desirable drug-like properties for potential use as PROTAC warheads.

**Figure 4.**
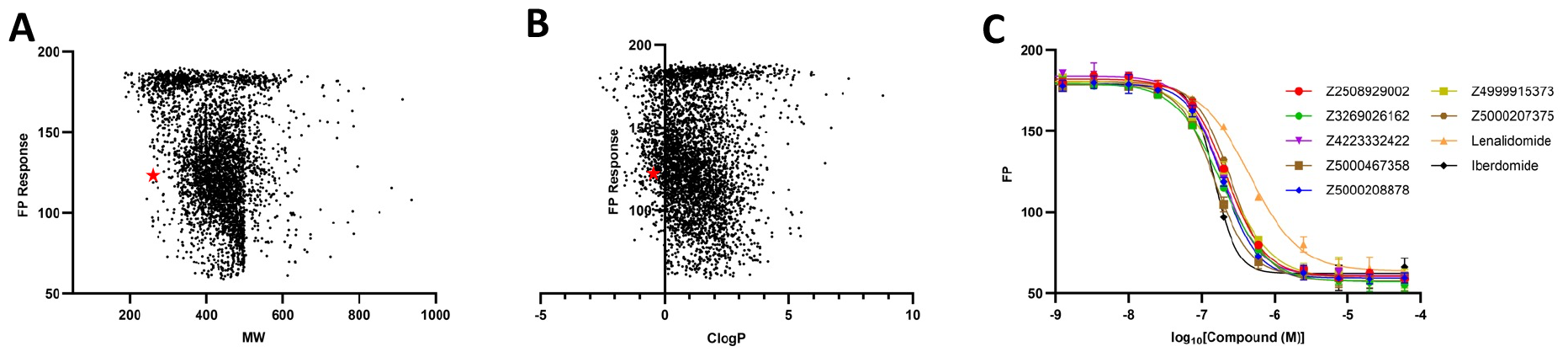
High affinity IMiD binders with favorable drug like properties : **A)** FP response vs molecular weight. Lenalidomide is labelled with a red asterix **B)** FP response vs clogP. Lenalidomide is labelled with a red asterix **C)** Dose response fluorescence polarization curves showing most potent binders with favorable drug like properties.

As expected, a number of substitutions in the phthaloyl ring are tolerated or even enhance CRBN-binding, while at the same time any close derivatives with even minor changes in the glutarimide part have low or no activity. Remarkably, highly enriched amongst the highest affinity binders in the primary FP screen are a previously undescribed class of compounds with sulfonamide derivatization of carbon 5/6 in the phthaloyl ring (Figure 3B). In total 122 sulfonamides exhibited an FP response equivalent to, or better than, lenalidomide, including the simplest sulfonomide derivative Z5000146157 (Table 2). The EC_50_ values of the top 50 hits were confirmed by dose-response measurements to be in the same range as iberdomide and a representative subset of these are displayed (Figure 3D)(Table 1). Given the recent identification and prevalence of aryl sulfonamides in DCAF-mediated molecular glues^25^ it will be highly interesting to explore this family of IMiD-based sulfonamide scaffolds for CRBN-based molecular glue-like functions as well as an approach to design PROTACS with sulfonamide containing linkers.

**Table 2:**
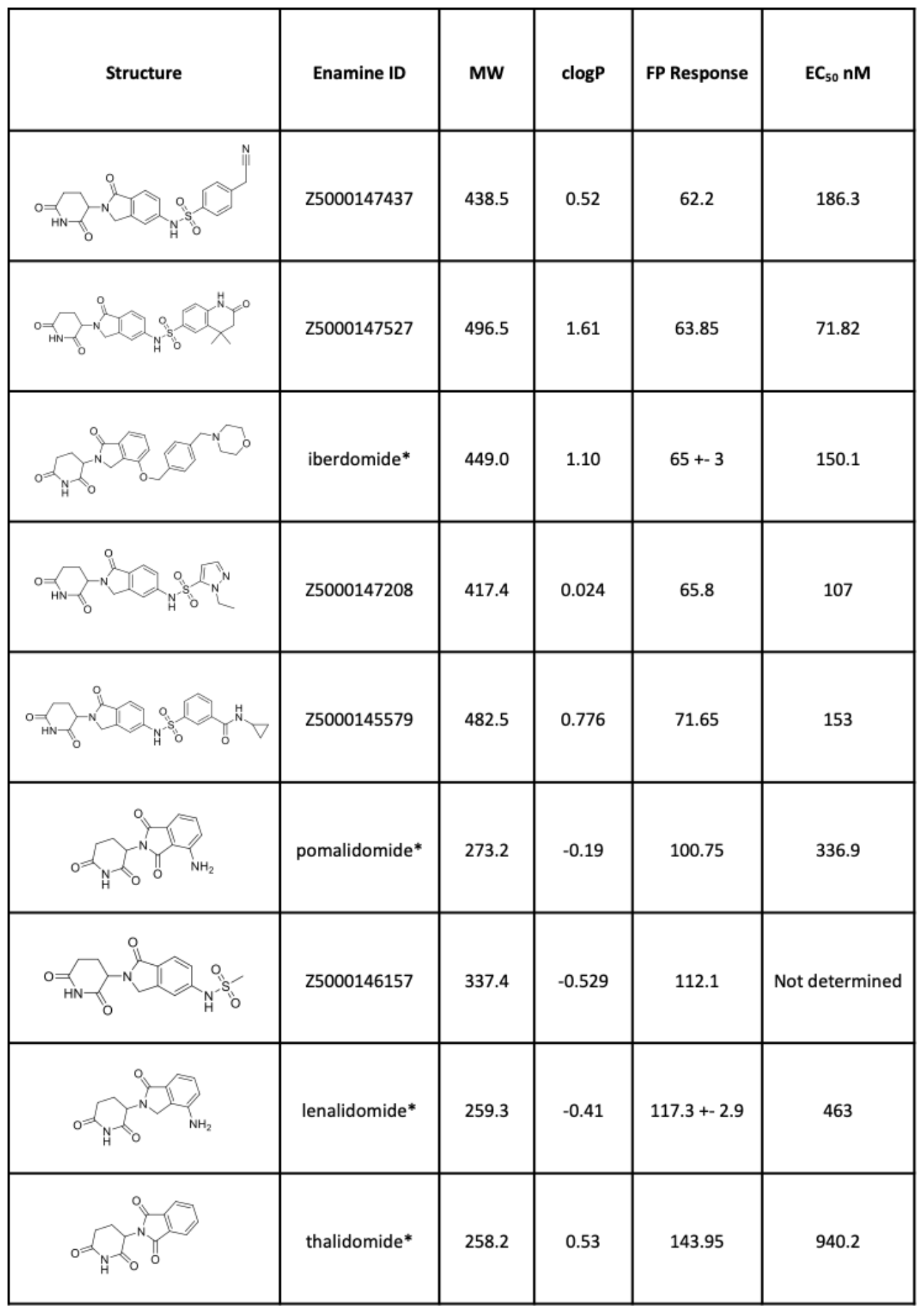
Example subset of sulfonamide based CRBN binders. *=control compounds

| Structure | Enamine ID | MW | clogP | FP Response | EC<sub>50</sub> nM |
| --- | --- | --- | --- | --- | --- |
|  | Z5000147437 | 438.5 | 0.52 | 62.2 | 186.3 |
|  | Z5000147527 | 496.5 | 1.61 | 63.85 | 71.82 |
|  | iberdomide* | 449.0 | 1.10 | 65 +- 3 | 150.1 |
|  | Z5000147208 | 417.4 | 0.024 | 65.8 | 107 |
|  | Z5000145579 | 482.5 | 0.776 | 71.65 | 153 |
|  | pomalidomide* | 273.2 | -0.19 | 100.75 | 336.9 |
|  | Z5000146157 | 337.4 | -0.529 | 112.1 | Not determined |
|  | lenalidomide* | 259.3 | -0.41 | 117.3 +- 2.9 | 463 |
|  | thalidomide* | 258.2 | 0.53 | 143.95 | 940.2 |

**Table 3:**
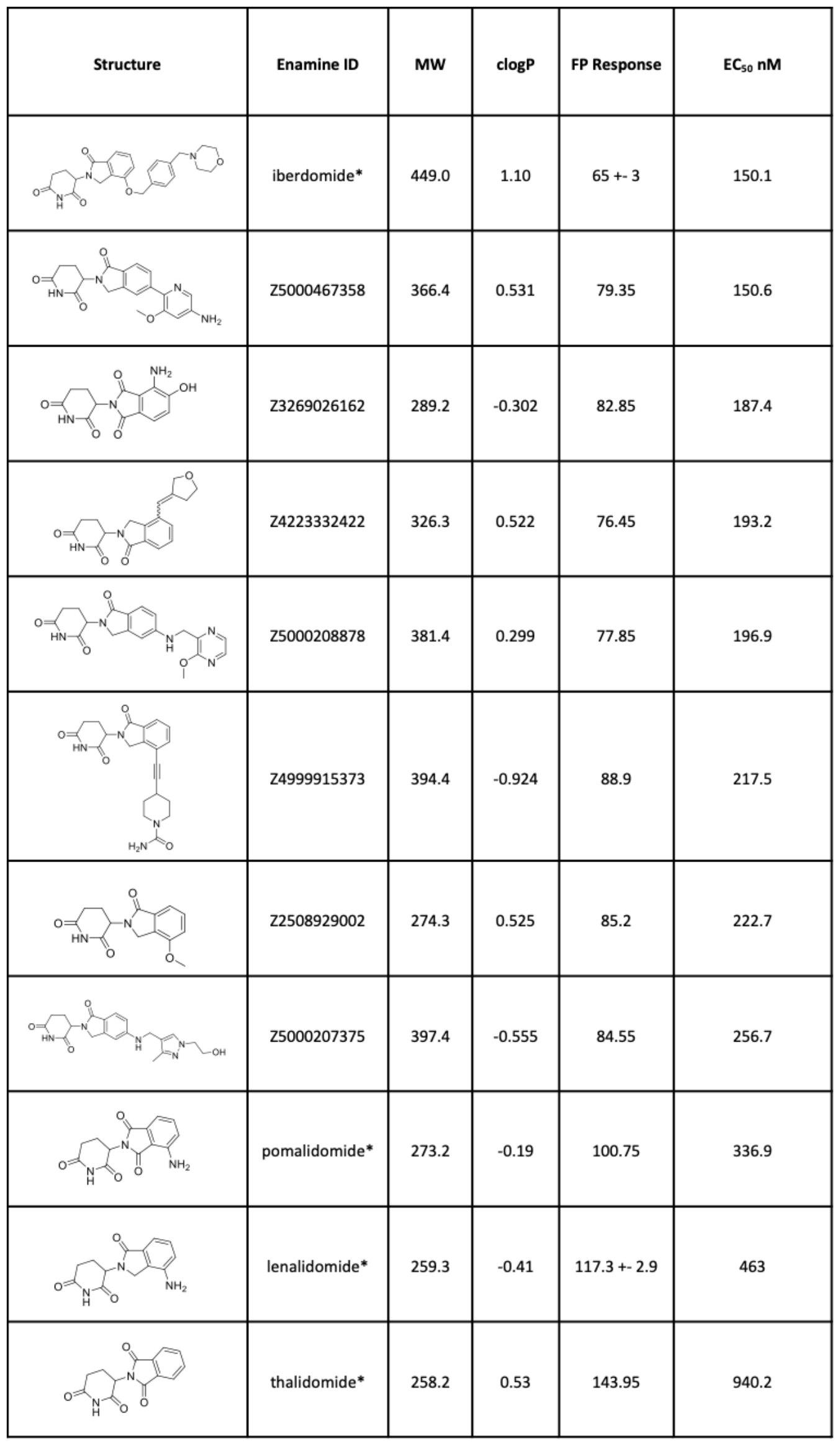
High affinity IMiD based binders with favorable drug like properties. *=control compounds

| Structure | Enamine ID | MW | clogP | FP Response | EC<sub>50</sub> nM |
| --- | --- | --- | --- | --- | --- |
|  | iberdomide* | 449.0 | 1.10 | 65 +- 3 | 150.1 |
|  | Z5000467358 | 366.4 | 0.531 | 79.35 | 150.6 |
|  | Z3269026162 | 289.2 | -0.302 | 82.85 | 187.4 |
|  | Z4223332422 | 326.3 | 0.522 | 76.45 | 193.2 |
|  | Z5000208878 | 381.4 | 0.299 | 77.85 | 196.9 |
|  | Z4999915373 | 394.4 | -0.924 | 88.9 | 217.5 |
|  | Z2508929002 | 274.3 | 0.525 | 85.2 | 222.7 |
|  | Z5000207375 | 397.4 | -0.555 | 84.55 | 256.7 |
|  | pomalidomide* | 273.2 | -0.19 | 100.75 | 336.9 |
|  | lenalidomide* | 259.3 | -0.41 | 117.3 +- 2.9 | 463 |
|  | thalidomide* | 258.2 | 0.53 | 143.95 | 940.2 |

## Discussion

In order to resolve the current limitation of CRBN expression for compound screening assays we developed an intermediate CRBN construct CRBN_ΔHBD that has been successfully designed for simple and more efficient production in *E.coli* expression systems. CRBN_ΔHBD has equivalent activity to CRBN:DDB1 towards thalidomide-based tool compounds, unlike previous *E.coli*-expressed CRBN constructs. Future use of CRBN_ΔHBD will therefore help circumvent the challenges of working with the heterogenous DDB1:CRBN complex in costly insect cell expression systems, providing a more effective construct for validating and ranking degrader compounds based on biophysical properties of binary and ternary complex formation. This data, obtained from the assays optimized here, will be highly desirable for integration with compound cellular target assays and degradation profiles for the rational design and optimization of PROTAC molecules and validation of MGs discovered through phenotypic cellular screening approaches.

The feasibility of the new construct in a high throughput screening approach was further tested by coupling the optimized binding assays with a CRBN-focused IMiD and diversity library. The library was specifically designed and prepared by Enamine to search both modified chemical scaffolds that may balance preferable drug like properties with improved ligand affinity and also sample diverse chemical space of the core IMiD scaffold for potential molecular glue-like functions. Enriched within the highest affinity hits in the library, we discover a previously unpublished sulfonamide derivatization of carbon 5/6 in the phthaloyl ring as a class of CRBN binders. It will be highly interesting to explore this family of IMiD-based sulfonamide scaffolds for CRBN-based molecular glue-like function given the recent discovery of aryl sulfonamide groups acting as molecular glues for other DCAFs^25^. We also identify the very recently published indazole scaffold with highly favorable drug like properties and potency of the minimal scaffold in the same range as pomalidomide. We further show binding potency can be improved with derivatization of the core scaffold. In addition to the indazole scaffold, we identify many further examples of the scaffolds that bind CRBN with higher potency, in the range of iberdomide (also known as CC-220), but without compromising molecular weight, solubility and other classical drug-like properties, allowing them to serve as potential new PROTAC handles.

The diversity of the chemical ligands uncovered within the screen should also provide potential for exploration of reduced off target toxicity. The potential teratogenicity of many of these new compounds is so far uncharacterized, but may also aid in the discovery of new, safer, next generation CRBN-based degraders. Together the protein and assays optimized here coupled with the new discovery of diverse IMiD based ligands will help shape future exploration of CRBN degradation space.

## Acknowledgements

We thank all the members of the Dikic laboratory and the members of the PROXIDRUGs consortium for their support and constructive discussion. We thank Sonja Welsch and Susann Kaltwasser from the Central Electron Microscopy facility of the Max Planck Institute of Biophysics for electron microscopy support. Furthermore, we acknowledge Proteros Biostructures GmbH (Munich, Germany) for protein supply of CRBN:DDB1 and BRD4 and thank Ana-Maria Esteves at the iBET-(Oeiras, Portugal) for the supply of CRBN TBD. Finally, we thank members of the SGC Frankfurt screening lab Benedict-Tilman Berger and Lewis Elson for technical screening support.

PROXIDRUGS as part of the initiative “Clusters4Future” is funded by the Federal Ministry of Education and Research BMBF (03ZU1109FA)

## Contributions

The present study was conceived by I.D., F.S., and H.B.. The experiments and analysis were performed by J.E., H.B. and J.V.. J.E. and H.B. carried out the majority of structural and biophysical experiments with technical support from E.L. and expertise from A.W., F.S., and I.D.. I.K. T.M. N.T. designed and selected the Enamine library. H.B. carried out primary screening and anaylsis with expertise from I.K. T.M. N.T.. J.V. carried out crosslinking mass spectrometry experiments with supervision and expertise of J.L.. H.B., J.E., F.S., and I.D. wrote the manuscript with contributions from all authors.

## Methods

### Cloning, Expression and Purification

Human CRBN-TBD (amino acids 318-426) was expressed and purified from *E. coli* essentially as described^26^Human BRD4-BD1(42-168) was expressed and purified as described^27^ Human CRBN:DDB1 complex (CRBN(40-442) and DDB1(1-1140)) was provided by Proteros Biostructures GmbH. For preparation of CRBN_ΔHBD, DNA insert containing CRBN_ΔHBD was cloned into pnic-MBP vectors, The plasmids were transformed into BL21(DE3) competent cells and single-colony inoculations were induced at an OD600 of 0.5– 0.7 with 0.2 mM IPTG for 16–18 h at 18 °C. Cultures were centrifuged and cell pellets stored at −80 °C. All cells were lysed by sonication and centrifuged at 35,000×g. The clarified cell extract was incubated with 2.5 mL of Ni-NTA resin pre-equilibrated with lysis buffer (50 mM HEPES pH 7.5, 500 mM NaCl, 10 mM imidazole, 5% Glycerol, 0.5 mM TCEP). The column was washed with 100 mL Binding Buffer (50 mM HEPES pH 7.5, 500 mM NaCl, 5% glycerol, 10 mM imidazole, 0.5 mM TCEP), 50 mL wash buffer (50 mM HEPES pH 7.5, 500 mM NaCl, 5% glycerol, 40 mM imidazole, 0.5 mM TCEP) and eluted with 15 mL of Elution Buffer (50 mM HEPES pH 7.5, 500 mM NaCl, 5% glycerol, 250 mM imidazole, 0.5 mM TCEP). The eluant fractions were concentrated to 5 mL and applied to a Superdex 200 16/60 column pre-equilibrated in GF Buffer (50 mM HEPES pH 7.5, 200 mM NaCl, 0.5 mM TCEP, 5% glycerol). Eluted protein fractions were pooled and concentrated to 10–17 mg mL−1. His6-tag removal was carried out by overnight treatment with TEV protease. Cleaved protein was further purified using a 5 mL HiTrap Q HP anion exchange chromatography column using a gradient of 25 mM NaCl to 500 mM NaCl across 15 column volumes (final buffer of eluent 50 mM HEPES pH 7.5, 200 mM NaCl, 0.5 mM TCEP, 5% glycerol).

### Fluorescence Polarization

Cy5-conjugated thalidomide (20 nM) was mixed with increasing concentration of either CRBN_DDB1, CRBN_ΔHBD or CRBN_TBD in 384-well microplates (Greiner) and incubated for 15 min at RT (50 mM HEPES pH 7.5, 200 mM NaCl, 0.5 mM TCEP, 5% glycerol 0.01 % Tween). The change in fluorescence polarization was monitored using a PHERAstar FS microplate reader (BMG Labtech) The Cy5-conjugated thalidomide bound fraction was calculated and the Ki was obtained from 2 technical and 2 biological duplicates. Compounds in Cy5-conjugated thalidomide displacement assay were dispensed in a 384-well microplate (Greiner) Echo-Dispenser normalized to 0.6% DMSO. Cy5-conjugated thalidomide (20 nM) and concentration of CRBN_DDB1, CRBN_ΔHBD or CRBN_TBD at 65% saturation in the K_D_ curves in 25 mM Hepes pH 7.5, 200 mM NaCl, 0.01% TWEEN. were performed. The change in fluorescence polarization across compound titrations was monitored using a PHERAstar FS microplate reader (BMG Labtech). Data from three independent replicates (n=3) were plotted and KI values obtained. CRBN_ΔHBD was screened using the same set up against a CRBN-focused library of 4480 compounds in duplicate at 500 nM concentration. Hits thresholds where selected 3 time beyond the standard deviation of DMSO control. Compound titration for calculation of EC_50_ of 175 hits were performed in duplicate.

### Thermal Stability

NanoDSF measurements were performed using Prometheus NT.Plex from NanoTemper. CRBN_ΔHBD at a final concentration of 0.3 mg/mL (50 mM HEPES pH 7.5, 200 mM NaCl, 0.5 mM TCEP, 5% glycerol, 1 % DMSO) was measured for stability changes in presence and absence of 50 µM ligands. The temperature was increased from room temperature to 95 °C, with a ramp of 1 °C per minute and stability monitored by changes in 330 fluorescence. All binding and control measurements were done in biological duplicates.

### Spectral Shift

Spectral shift measurements were performed using the Monolith X (NanoTemper). CRBN protein was labelled with RED-Maleimide 2nd Generation dye (NanoTemper) optimized for degree-of-labeling of 1:1 molar ratio calculated by nanodrop. An 11-point PROTAC dilution series from 60 µM to 1.25 nM, normalized to 0.6 % DMSO plus one DMSO control, were dispensed into 10 uL of Labelled CRBN_ΔHBD 10 mM concentration (20 mM HEPES, 200 mM NaCl, 1 mMT TCEP, 0.01% Tween 20, 5% Glycerol, pH 7.5) in the presence and absence of purified BRD4 at a fixed concentration of 5 µM. The samples were loaded to Monolith Premium Capillaries (NanoTemper), after 5 min centrifugation of the plate at 3,000 x g at 25 °C. The measurements were performed with medium IR power and auto-excitation power at 25 °C and fluorescence was detected and plotted for ratio metric changes in 650 and 670 nm wavelengths. Data plotting, K_D_ calculation was carried out using MO.Control software (NanoTemper). All measurements were performed in 2 technical and 2 biological duplicates.

### Mass Photometry

Mass photometry measurements were performed using a Two MP Auto (Refeyn). Zeiss high-precision microscope cover glasses 24 x 50 mm with CultureWell reusable gaskets (50 - 3 mm, DIA x 1 mm depth, 3 - 10 µL) were transferred to the laser lens covered with a drop of Immersol 518 F from Zeiss. 16 µL of mass photometry buffer (20 mM HEPES, 150 mM NaCl, pH 7.5) followed by 4 µL of 100 nM protein solution was transferred to a gasket well. To reach the final assay concentration 20 nM. The ratiometric contrast was evaluated using MP Discovery software in a 1 min movie and converted to mass values, using a standard curve of marker proteins of known mass.

### Analytical SEC

Analytical SEC was carried out using a 1260 Infinity II high pressure liquid chromatography (HPLC) system from Agilent Technologies (Santa Clara, USA), with an AdvanceBio SEC 300 Å 2.7 µm column. The final assay concentration of the proteins was set to 12.5 µM in a total volume of 35 µL (20 mM HEPES, 200 mM NaCl, 1 mM TCEP, 0.01% Tween 20, 5% Glycerol, pH 7.5). Ligands and DMSO controls were pre-incubated for 10 min at RT followed by centrifugation for 15 min at 22,000 x g. The samples were transferred to a 96-well plate and stored at 15 °C in the HPLC autosampler. The proteins were detected at a wavelength of 280 nm and complex formation was determined according to retention time and integral of the peaks.

### X-linking Mass Spectrometry

Proteins at 15 µM were preincubated for 15 mins at RT in presence and absence of DBET6 PROTAC at 1:1 or 1:2 molar ratio (50 mM HEPES pH 7.5, 200 mM NaCl, 0.5 mM TCEP, 5% glycerol). Disuccinimidyl suberate crosslinker (ThermoFischer) was spiked 3 times over 30 mins to final conc of 0.25 mM and crosslinking reaction was quenched by addition of 150 mM ammonium bicarbonate. The gel band corresponding to the crosslinked complex was excised and digested with trypsin. Excised gel bands were digested using the In-Gel Tryptic Digestion Kit (Thermo Fisher Scientific, Dreieich, Germany). The gel pieces were destained, reduced and alkylated according to the manufacturers protocol. Digestion was performed overnight at 30 °C in a 25:1 ratio of protein:trypsin (SERVA, Heidelberg, Germany). Digested peptides were transferred into a clean tube with 5 µl 0.1% formic acid (FA) in H2O. Further peptides were extracted from the gel pieces using 10 µl of 1% FA in H2O and added to the digestion mixture. Peptides were injected onto an Acclaim PepMap C18 capillary trapping column (particle size 3 µm, L = 20 mm) and separated on a ReproSil C18-PepSep analytical column (particle size = 1,9 µm, ID = 75 µm, L = 15 cm, Bruker Coorporation, Billerica, USA) using a nano-HPLC (Dionex U3000 RSLCnano) at a temperature of 55 °C. Trapping was performed at a flow rate of 6 μL/min for 6 minutes using a loading buffer composed of 0.05% trifluoroacetic acid in H2O. Peptide separation was carried out at a constant flow rate of 400 nL/min using a gradient of water (buffer A: 100% H2O and 0.1% FA) and acetonitrile (buffer B: 80% ACN, 20% H2O, and 0.1% FA). The gradient increases from 4% to 48% buffer B in 30 min. All solvents were LC-MS grade and purchased from Riedel-de Häen/Honeywell (Seelze, Germany). Eluting peptides were analyzed in data-dependent acquisition mode on an Orbitrap Eclipse mass spectrometer (Thermo Fisher Scientific, Dreieich, Germany) that is connected to the nano-HPLC by a Nano Flex ESI source. MS1 full scans were acquired over a range of 380–1400 mass-to-charge ratio (m/z) in the Orbitrap detector (resolution = 60k, automatic gain control (AGC) = 4e6, and maximum injection time: 50 ms). Sequence information was acquired by a “ddMS2 OT HCD” MS2 method with a fixed cycle time of 2 s for MS/MS scans. MS2 scans were generated from the most abundant precursors with a minimum intensity of 5e4 and charge states from 3 to 8. Selected precursors were isolated in the quadrupole using a 1.6 Da window and fragmented using higher-energy collisional dissociation (HCD) at 25.3% normalized collision energy. For Orbitrap MS2, an AGC of 1e5 and a maximum injection time of 70 ms were used (resolution = 30k). Dynamic exclusion was set to 60 s with a mass tolerance of 10 parts per million (ppm). The sample was measured in technical duplicates. MS raw data were processed using the MaxQuant software (v2.1.4.0) with customized parameters for the Andromeda search engine. Spectra were matched to a FASTA file containing the sequences of human CRBN and BRD4 (UniProtKB, March 2023) and a decoy and contaminant database. In addition, a list of E. coli proteins that were identified with > 5 unique peptides in the sample on a previous search against the E. coli proteome (UniProt Proteomes, March 2023) was added. The minimum tryptic peptide length was set to 7 with a maximum of two missed cleavage sites. Carboxyamidomethylation of cysteine residues (static modification), methionine oxidation and acetylation of the protein N-terminus were included (variable modifications) in the search parameters. Precursor mass tolerance was set to 4.5 ppm and fragment ion tolerance to 20 ppm. A false discovery rate (FDR) below 1% was applied for the identification of crosslinks, peptides, and modifications. Crosslinked peptides were manually inspected and evaluated for consistent identification in both technical replicates. All proteomics data (including acquisition and data analysis parameters) associated to this manuscript will be deposited at the ProteomeXchange Consortium (http://proteomecentral.proteomexchange.org).

### Negative Stain Electron Microscopy

CRBN_ΔHBD at 0.15 mg/ml (150 mM NaCl, 25 mM HEPES pH 7.5, 0.5 mM TCEP) was pre-incubated with Homo-PROTAC at 1:2 molar ratio (1 % DMSO). The samples were diluted 10 fold immediately prior to grid staining Carbon coated grids (C flatTM, 300 Mesh) from were glow discharged. 3 µL of the protein sample was transferred to the grid and incubated for 30 s. negatively stained with 2% (w/v) uranyl formate and analyzed by negative-stain electron microscopy with a Rio16 CMOS camera (Gatan) on a Tecnai Spirit (FEI Company) transmission electron microscope (TEM) operated at 120 kV.

### CRBN-focused library

All compounds for the CRBN-focused library were obtained at Enamine using in-house developed procedures. 4480 compounds (10 µL of 10 mM DMSO solutions) were delivered in 384-well echo plates (14 plates), prepared using Labcyte #LP-0200, (320 compounds per plate, first two and last two columns empty).

## Supplementary Figures

**Supplementary Figure 1:**
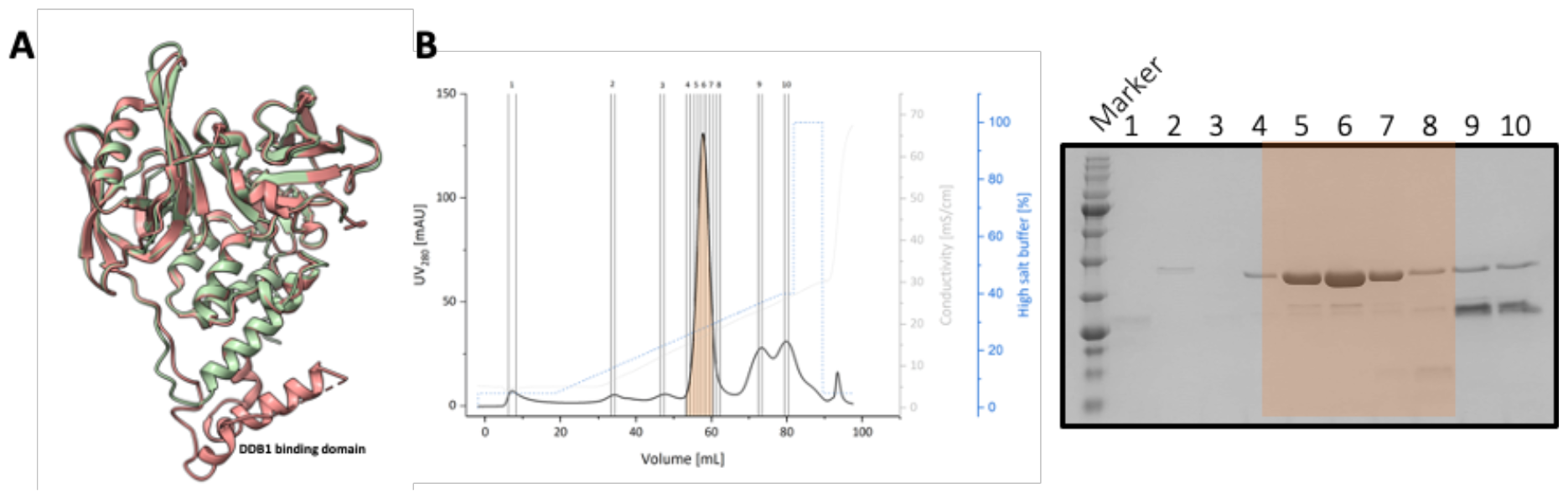
Alpha fold analysis and purification of CRBN_ΔHBD. **A)** Overlay of alpha fold generated structure of CRBN_ΔHBD (coloured green) with CRBN from PDB 6BOY^14^ (coloured red) **B)** Purification of CRBN_ΔHBD. Left shows anion exchange chromoatogram. Right shows SDS page gel with final purity of over 95 %.

**Supplementary Figure 2:**
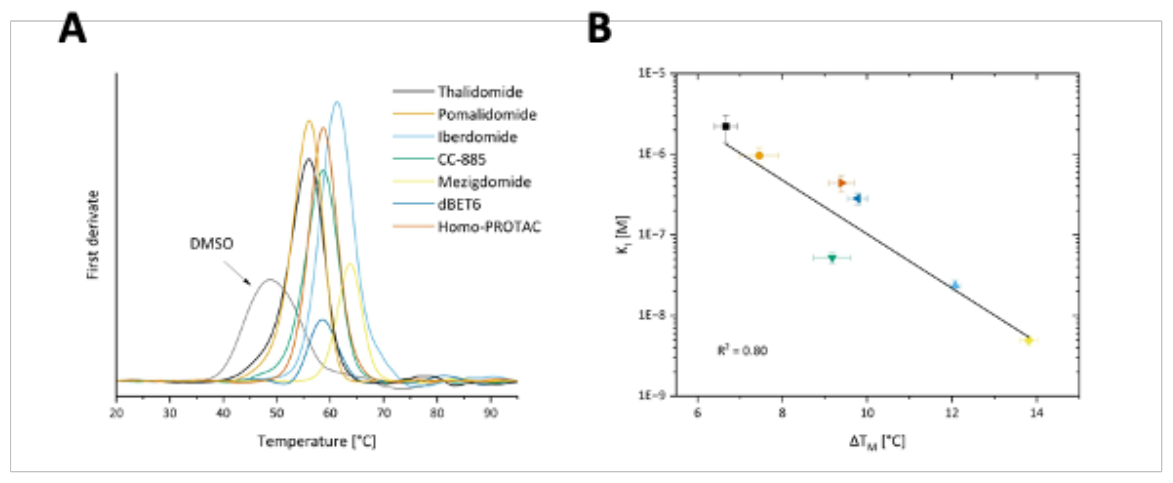
Thermal shift assay of tool CRBN binders: **A)** Thermal shif assay showing stability shifts of CRBN_ΔHBD in the presence of tool compounds **B)** Correlation plotting relationship between thermal stability shift and K_i_ of tool molecular glues.

**Supplementary figure 3:**
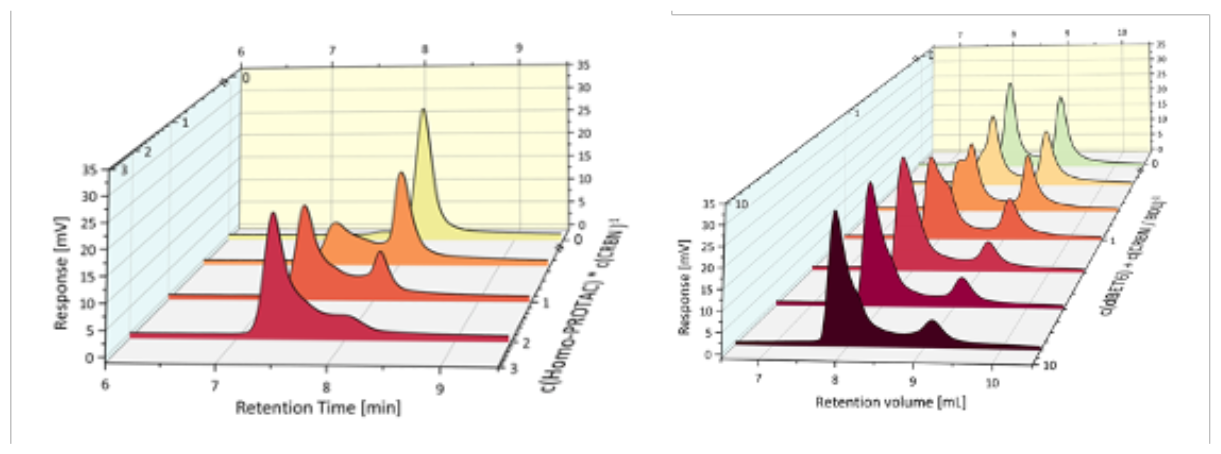
Size exclusion chromatography showing ternary complex formation of CRBN_ΔHBD: Left shows changes in retentiaon volume of CRBN_ΔHBD with titration of homo-PROTAC. Right shows changes in retentiaon volume of CRBN_ΔHBD and BRD4-BD1 with titration of DBET6.

**Supplementary figure 4:**
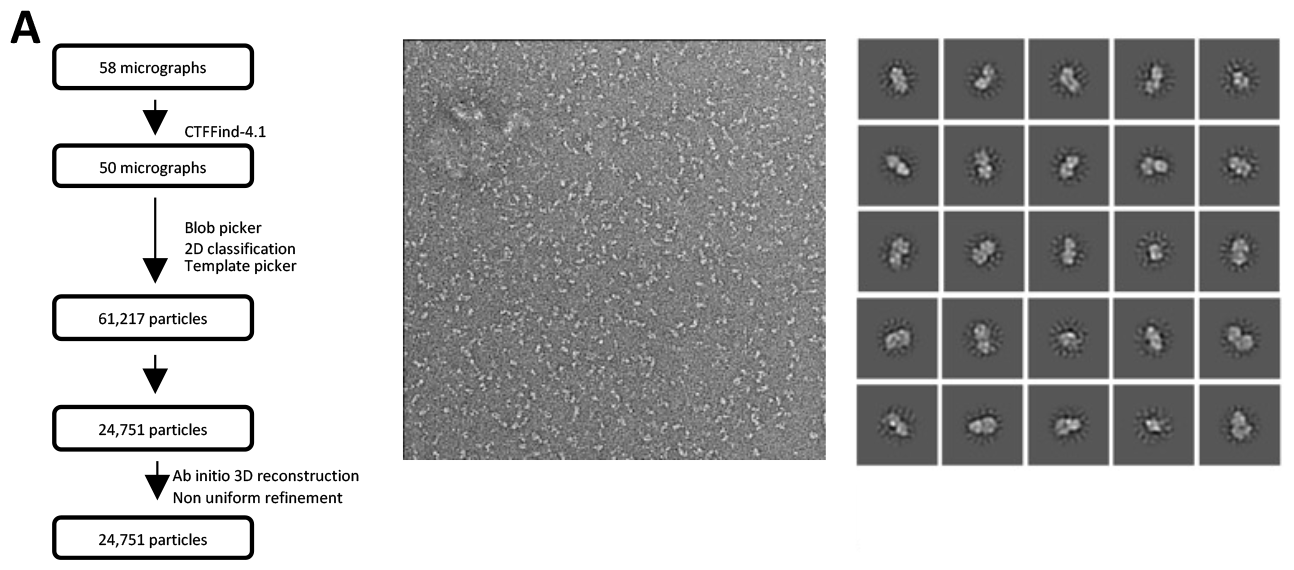
Negative stain microscopy processing: Workflow, example micrograph and 2D classes of CRBN_ΔHBD in presence of homo-PROTAC.

**Supplementary Figure 5:**
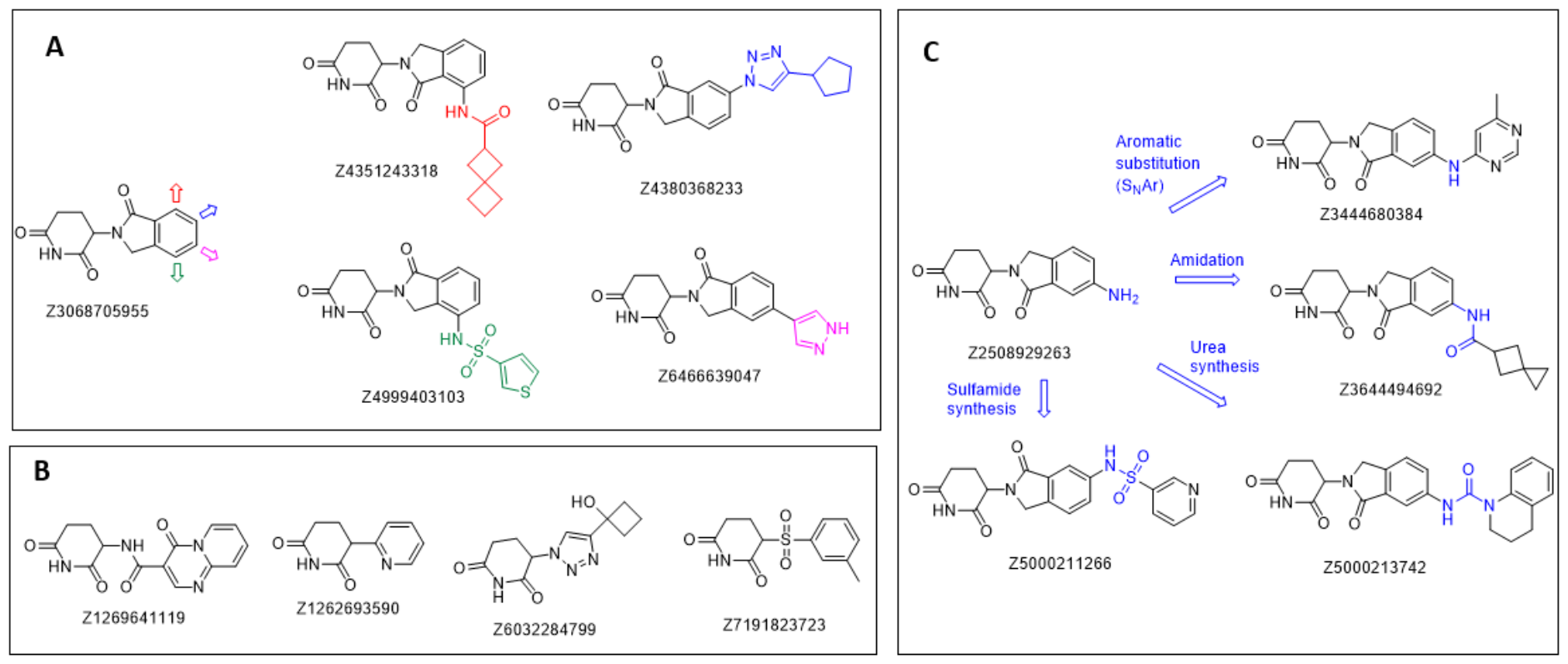
Design of CRNB-focused library: A) Example of compound selection with diverse substitution of the core in different positions of benzene ring. B) Exanples of compounds with variation in the core structure. C) Example of synthesis of diverse compounds for the library based on core Building Blocks

**Supplementary figure 6:**
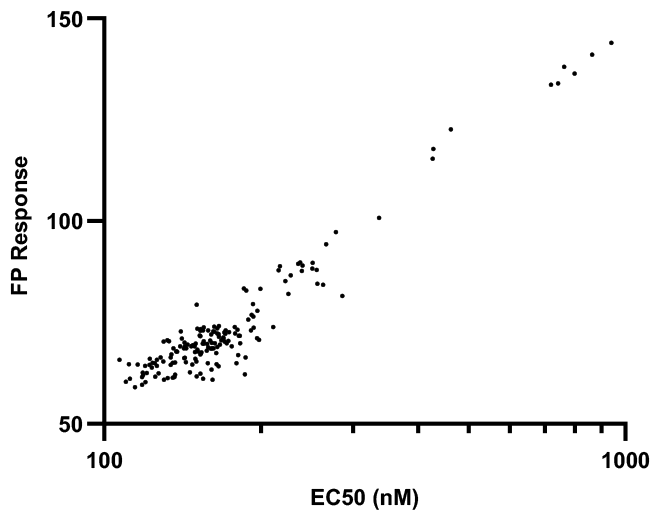
EC_50_ vs Primary FP screen response.

**Supplementary figure 7:**
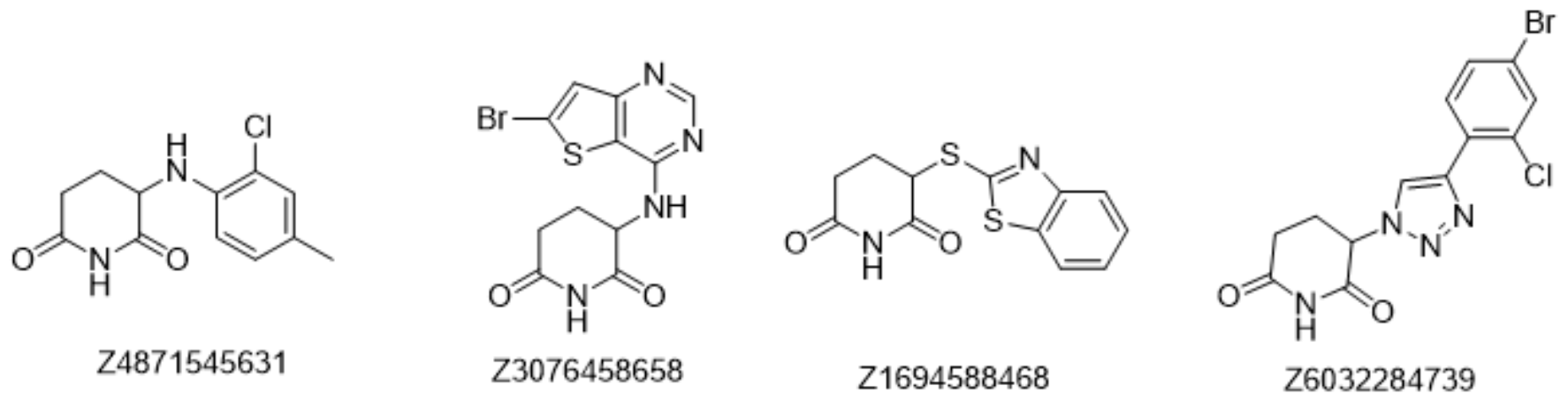
Examples of identified weaker binders with non-phthaloyl containing scaffolds.

**Supplementary Table 1:**
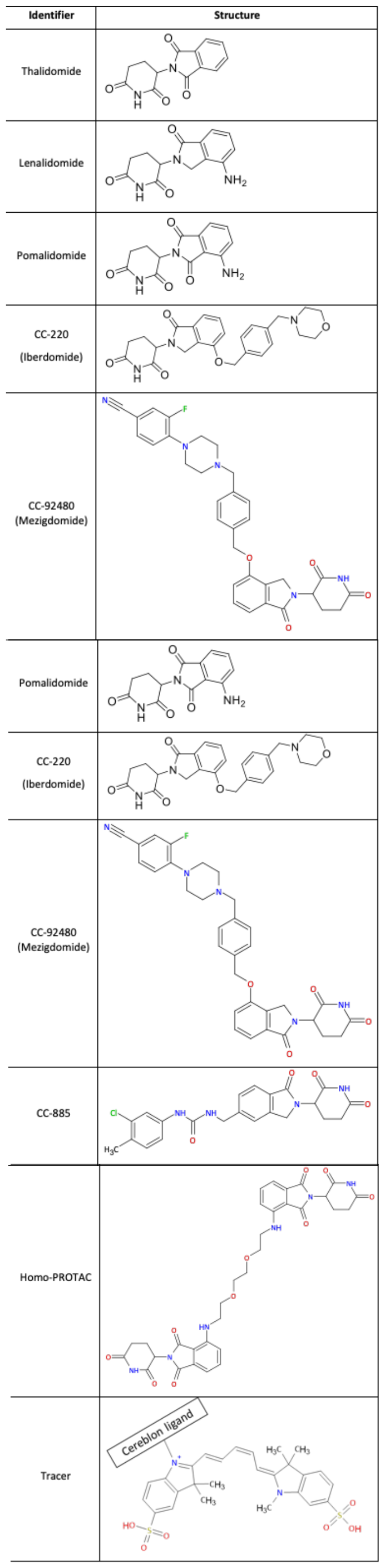
Tool compounds used in this study. All ligands were provided by Merck Healthcare KGaA

| Identifier | Structure |
| --- | --- |
| Thalidomide |  |
| Lenalidomide |  |
| Pomalidomide |  |
| CC-220<br>(Iberdomide) |  |
| CC-92480<br>(Mezigdomide) |  |
| Pomalidomide |  |
| CC-220<br>(Iberdomide) |  |
| CC-92480<br>(Mezigdomide) |  |
| CC-885 |  |
| Homo-PROTAC |  |
| Tracer |  |

## Notes

### Competing Interest Statement

The authors have declared no competing interest.

